# CRISPR herd immunity against a transducing phage underlies adaptive homologous recombination in resting bacteria

**DOI:** 10.1101/2025.10.01.679802

**Authors:** P. Payne, P. Plevka, J. P. Bollback, C. C. Guet

**Affiliations:** Institute of Science and Technology Austria; Klosterneuburg, Austria; Institute of Infection, Veterinary and Ecological Sciences, University of Liverpool; Liverpool, United Kingdom; Central European Institute of Technology, Masaryk University; Brno, Czech Republic

## Abstract

Homologous recombination (HR) is ubiquitous across evolution, driving adaptation by reshuffling standing genetic variation. Although bacteria lack meiotic recombination, HR extensively shapes their genomes. However, the mechanisms and ecological conditions sustaining frequent HR in bacteria remain unclear. Using *Escherichia coli*, we reveal how frequent recombination emerges from herd immunity to a generalized transducing phage. Herd immunity–established here via CRISPR immunity–maintains genetic polymorphism and enables stable host–phage coexistence, thereby promoting genome-wide gene flow and accelerating adaptation through recombination up to two orders of magnitude relative to *de novo* mutations. Notably, we show that recombination occurs in stationary phase and is mediated by RecG, which has been previously reported to be regulated by the stringent response – a bacterial reaction to nutrient deprivation and other stress conditions. Bacterial herd immunity thus fulfills an unexpected role of promoting adaptation by HR. This mechanism helps explain the enigmatic high rates of HR across bacterial populations, clarifies how bacteria adapt as resources wane, and suggests a broader evolutionary role for bacterial immune systems beyond individual defense.

## Introduction

Homologous recombination is a pervasive evolutionary process that can accelerate adaptation of populations by reshuffling standing genetic variation: it can bring beneficial mutations that are present at different genomic loci in different individuals into the same genomic background (*1, 2*). Recombination also helps reduce the accumulation of deleterious mutations, which can otherwise lead to population decline or extinction through an evolutionary process known as Muller’s ratchet (*3*). In particular, recombination was shown to be beneficial in changing environments (*4, 5*). This is best exemplified in unicellular eukaryotes, which reproduce clonally when nutrients are abundant, but use independently evolved starvation-induced sexual reproduction in response to resource scarcity (*6*–*9*). From the genomic point of view, frequent HR results in an unmistakable signature: near-random associations among mutations at different loci across a population (*10*).

Although bacteria lack meiotic recombination, they often display population genomic signals demonstrating frequent HR: near-random associations among loci and high *r/m* ratio (recombination rate to mutation rate) (*11*–*15*). HR is often strongly biased toward within-population exchange; combined with spatial niche structuring, this bias can impose bacterial species cohesiveness, strong reproductive isolation between species and even promote ecological and genetic divergence (*16*–*18*).

It is well known that bacteria can exchange genetic information by horizontal gene transfer (HGT) (*19*). Yet the ecological and mechanistic bases sustaining such high HR in natural bacterial populations remain unclear (*12, 18*). Canonical HGT routes are well characterized *in vitro*, but each faces ecological constraints *in situ*. Natural transformation, which requires competence and uptake of extracellular DNA, has been demonstrated for only about 80 bacterial species and is typically transient and environmentally induced (*20*). Conjugation depends on direct cell-to-cell contact and maintenance of conjugative plasmids, and it is limited by spatial structure, plasmid burden, and incompatibility, and only rarely mediates chromosomal HR (*21*). Generalized transduction is an infrequent by-product of lytic infection. Most phage–host encounters end in lysis rather than host DNA transfer, yielding low transduction probabilities and lysis-dominated dynamics in nature (*22, 23*). Other proposed contributors, such as gene transfer agents (GTAs) and extracellular membrane vesicles, lack strong quantitative support for major population-level effects (*24, 25*). Taken together, none of the mechanisms listed above can reliably explain the high HR rates observed in natural bacterial populations

The rates of HGT can be influenced by bacterial defense (immune) systems, which are traditionally considered barriers to HGT because they can act against mobile genetic elements (*26*). However, recent comparative genomics analyses reveal that on evolutionary scales, many immune systems (restriction modification (RM) systems, AbiE, AbiJ, CBASS, Gabija, Wadjet, and others) are associated with elevated rates of gene gain, indicating that they could potentially facilitate HGT and HR (*27, 28*). Furthermore, one study found that CRISPR-Cas (<u>c</u>lustered <u>r</u>egularly interspaced <u>s</u>hort <u>p</u>alindromic <u>r</u>epeats – <u>C</u>RISPR <u>as</u>sociated genes) system in *Petrobacterium atrosepticum* can enhance generalized transduction by increasing survival rates of immune cells when high concentration of a phage lysate is applied to the cultures (*29*).

The question remains: how do bacterial immune systems modulate HR within populations?

Here, we present a mechanism that arises from molecular interactions between bacterial immunity and phage DNA and, at the population level, maintains genetic polymorphism and drives frequent homologous recombination in bacterial populations: herd immunity (Fig. 1A). When a fraction of the population carries an immune system that interferes with phage DNA, it effectively removes phages from the population and protects a phage-susceptible fraction of bacteria. This interaction imposes frequency-dependent selection on the immune strain, which can increase only up to the herd immunity threshold frequency (Fig. 1B). At the same time, the remaining phage-susceptible cells allow the phage to persist in the population at an equilibrium concentration. Consequently, if the phage is capable of generalized transduction, gene flow from the phage-susceptible to the immune fraction of the population can be established. Because herd immunity has already been demonstrated experimentally for CRISPR immunity (*30*), we set out to test this hypothesized mechanism using *E. coli* with engineered CRISPR immunity and a virulent variant of the generalized transducing phage P1.

**Figure 1.**
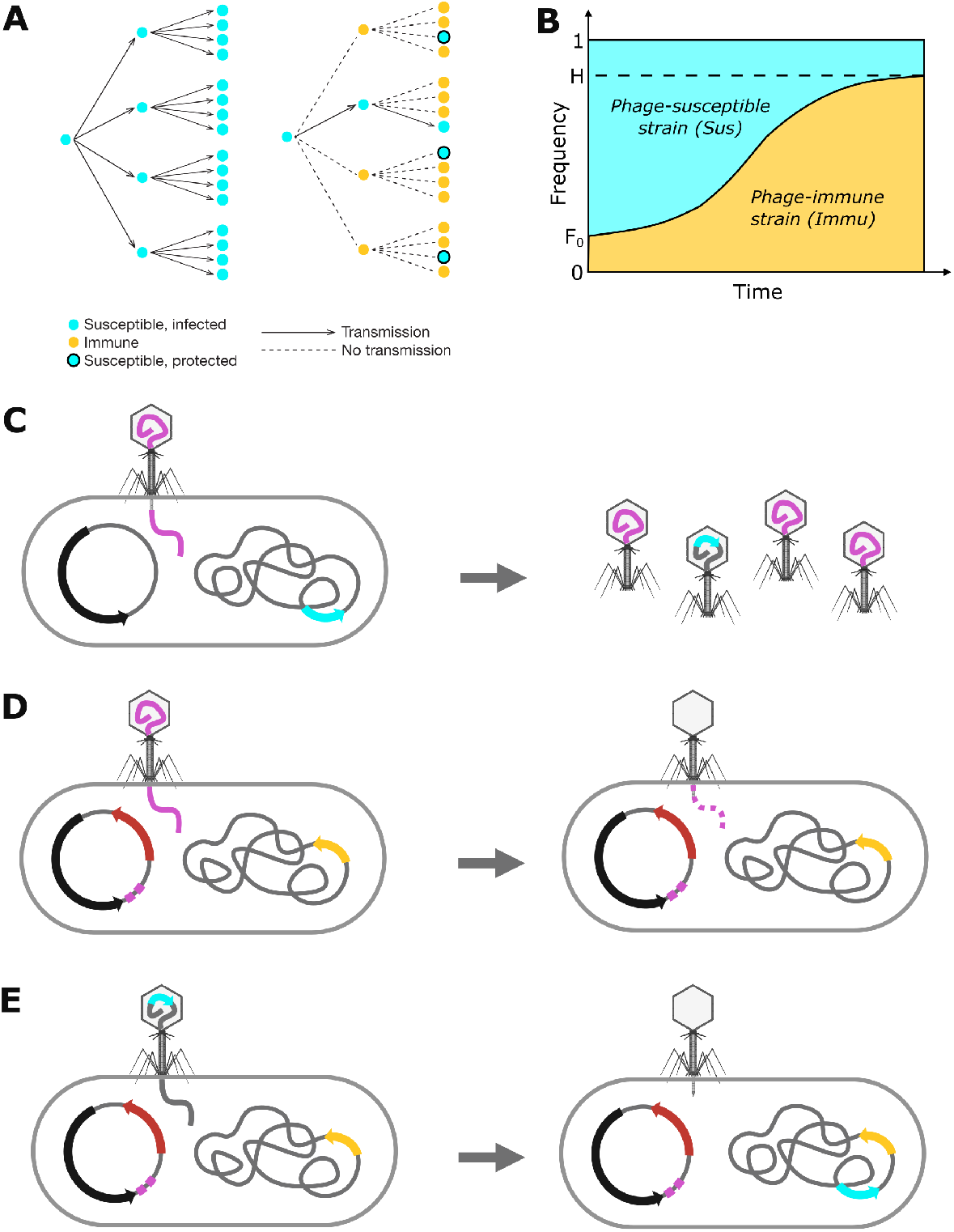
Herd immunity and recombination dynamics in the experimental system. **(A)** Illustration of principles of herd immunity. Without any immune individuals, each infected susceptible individual (cyan) infects (solid arrows) four more susceptible individuals. Thus, the pathogen spreads exponentially in the populations. With ¾ of immune individuals (yellow) in the population, transmission of the pathogen is blocked (dashed lines) and some susceptible individuals are protected by herd immunity (cyan in black circles). **(B)** Herd immunity-induced frequency-dependent selection on the immune strain (*Immu*). Starting from an initial frequency F_0_, *Immu* can only reach the herd immunity threshold frequency *H*, maintaining the susceptible strain, *Sus*, in the population. **(C)** *Sus*, which carries a non-targeting control plasmid encoding Cas9 (black arrow) with no spacers and a chromosomal cassette comprised of *cfp* and spectinomycin resistance genes (*cfp*-*specR*, cyan arrow), is being infected by a wild type P1 phage (left), giving rise to phage progeny with a fraction of them carrying the chromosomal insertion cassette (right). (**D)** Immu carrying a plasmid with Cas9 (black arrow), two P1 phage targeting spacers (purple squares), *rfp* gene (red arrow) for red fluorescence, and a chromosomal cassette comprised of *yfp* and kanamycin resistance genes (*yfp-kanR*, yellow arrow) is being infected by the wild-type phage (left). Upon ejection of the wild-type phage DNA into Immu, the DNA is targeted and cleaved by the CRISPR-Cas9 system (right). (**E**) When a stretch of the Sus donor DNA containing the *cfp-specR* cassette is injected into Immu (left), the transduced DNA is left intact and can recombine into the host cell chromosome. This gives rise to a strain expressing both chromosomally encoded *cfp* and *yfp* casssettes, as well as the plasmid encoded *rfp* (right).

In particular, we focus on investigating (i) whether herd immunity-driven HR can outpace *de-novo* mutations and (ii) the conditions under which bacteria resort to recombination – is it associated with exponential growth or, as in unicellular eukaryotes, linked to stationary phase.

## Results

### Experimental system

We engineered two strains of *E. coli* MG1655. The first strain contained a chromosomal constitutively expressed cyan fluorescence protein coding gene coupled with spectinomycin resistance gene (*cfp-specR*) at the phage λ attachment site *attB* (Fig. 1C). The second strain contained a chromosomal constitutively expressed gene for yellow fluorescent protein coupled with kanamycin resistance gene (*yfp-kanR*) at the phage P21 *attB* site (Fig. 1D). Thus, each strain was resistant to one of the antibiotics and encoded one of the fluorescence proteins. Adaptation to the simultaneous presence of both antibiotics could occur either through a *de novo* mutation (at any site on the chromosome) conferring resistance to the other antibiotic, or by HR bringing the resistance-fluorescence cassette from the other strain. Therefore, recombination gave rise to colonies, which grew on both antibiotics and expressed both YFP and CFP, whereas *de novo* mutation led to colonies expressing either YFP or CFP, but not both. This experimental design allowed us to directly compare rates of adaptive *de novo* mutations and rates of HR in the absence of selection.

As hypothesized above, HR in the genetic system could occur due to the presence of CRISPR-based immunity to a phage capable of generalized transduction. As the endogenous *E. coli* CRISPR systems of strain MG1655 are not active under standard laboratory conditions (*31*), we provided the engineered *yfp-kanR* strain with a very low-copy plasmid containing the *cas9* gene from *Streptococcus pyogenes* and two spacers targeting the phage P1 genome (Fig. 1D, strain *Immu* hereafter), which provided immunity against the phage. Plasmid-based immunity mimics natural conditions, as CRISPR-Cas systems are widely distributed both on chromosomes and on plasmids (*32*).

In order to distinguish immune and phage-susceptible cells, we inserted a constitutively expressed red fluorescent protein coding gene (*rfp*) into the plasmid providing immunity against the P1 phage. The phage-susceptible strain, referred to as *Sus*, was transformed with the same low-copy-number plasmid containing *cas9*, but without *rfp* and the spacers targeting phage P1 (Fig. 1C). In a competition assay we verified that the growth rate of the *Sus* strain did not differ significantly from the *Immu* strain (Suppl. text, Fig. S1).

In our experimental system, HR is enabled by the following processes: Upon multiplication of the wild-type P1 phage on the *Sus* strain, heterogeneous progeny is produced with a fraction of transducing phages carrying about 100kb long fragments of the strain *Sus* bacterial chromosomal DNA (Fig. 1C). When an *Immu* bacterium is infected by a wild-type infectious phage particle, it cleaves its DNA by the CRISPR-Cas immune system and survives (Fig. 1D). When the phage particle is transducing DNA, the DNA is left intact and can recombine into the *Immu* strain chromosome (Fig. 1E). Thus, an *Immu* bacterial cell can “explore” multiple phage particles and survive many wild-type phages before it encounters a transducing one. Simultaneously, the *Immu* strain acts as a sink for the phage, which leads to protection of the *Sus* strain by herd immunity.

### Recombination increases in resting bacteria

We started by testing cultures that grew exponentially and reached stationary phase during the experiment. We inoculated overnight cultures of the strains 1:100 in fresh growth medium in a ratio close to the expected herd immunity threshold (*30*), *i.e*., 95% *Immu*: 5% *Sus*, which we determined using a mathematical model of the experimental system (Suppl. text, Fig. S3, S4). We added the phage at an initial concentration of 1:10 relative to bacterial cells (multiplicity of infection – MOI 0.1). Then we incubated the cultures in a shaking aerated incubator for 24 hours and sampled at 3, 6, 12, 18, and 24 hours post-inoculation (hpi). The growth medium did not contain any spectinomycin or kanamycin, and thus there was no selection for the recombining loci and mutations during the experiment. Selection occurred only after sampling on plates containing antibiotics selecting for both resistance markers, enabling us to count colony forming units (CFUs) formed by bacteria that have either mutated or recombined the second selection marker into their genome.

We found that the concentration of CFUs surviving selection because of HR started exceeding the concentration of mutant CFUs at 6 hpi (Fig. 2A, B, solid vs. dashed lines). This indicates that adaptation by recombination is superior to adaptation by *de novo* mutations when bacteria begin to enter stationary phase (Fig. 2B). Interestingly, from this time point on, the number of recombinant CFUs kept increasing even though the total population of bacteria was slowly declining. This resulted in a rapid increase of the frequency of recombinants (Fig. 2C) independent of the bacterial population size. At the end of the experiment (24 hpi), the concentration of recombinants reached about 380-fold more than the concentration of spontaneous mutants, showing that in early stationary phase, recombination is far more important for adaptation than point mutations. The concentration of CFUs adapted by spontaneous mutations was relatively stable throughout the experiment (Fig. 2A, dashed line) with a slight increase at 6 and 12 hpi and a drop at 18 and 24 hpi. This observation is consistent with an expected mutation rate in the population (*33*) and the fact that *de novo* mutations occur during exponential growth. The drop at 18 and 24 hpi can be attributed to an expected fitness cost of the mutations for antibiotic resistance in the absence of antibiotic selection pressure.

**Figure 2.**
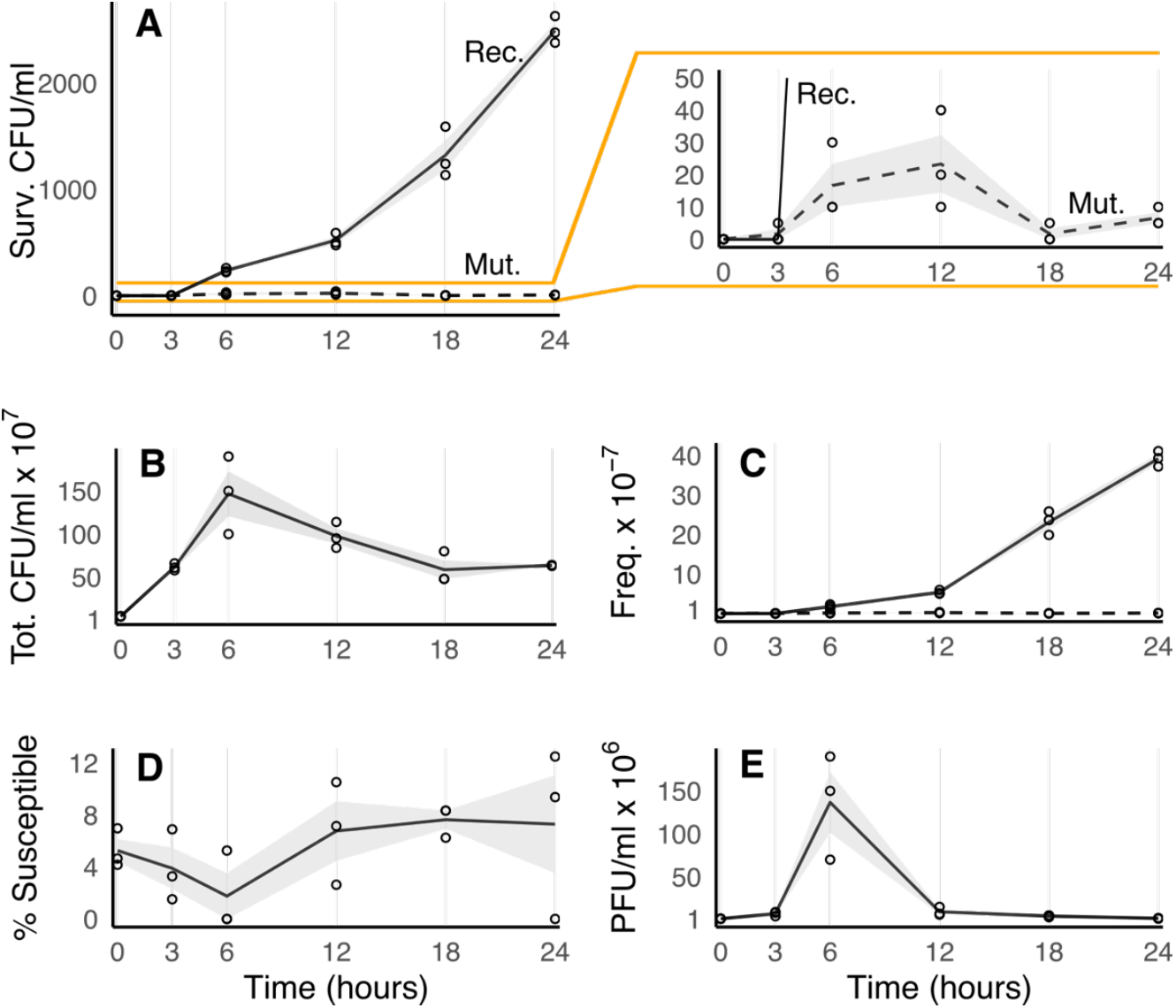
Recombination dynamics in initially exponentially growing cultures. **(A)** Left: Concentration of bacteria (colony forming units, CFUs), which survived kanamycin and spectinomycin selection on sampling plates due to recombination of chromosomally encoded cassettes (solid line, Rec.) and concentration of mutant CFUs with only one of the cassettes and a *de novo* mutation for resistance to the other antibiotic (dashed line, Mut.). Right: Zoom on mutant CFUs. (**B)** Concentration of total CFUs increases until 6 *hpi*, when the cultures reach stationary phase and their concentration starts slowly declining. **(C)** Frequencies of recombinant (solid line) and mutant (dashed line) CFUs. **(D)** Phage-susceptible cells remain in cultures throughout the experiment. **(E)** Concentration of viable phages (expressed in plaque forming units, PFUs) in experimental cultures peaks at 6 hpi, after which phage concentration converges to its equilibrium concentration around 10^6^ PFU/ml. In all panels the shaded areas around the curves show standard error of the mean.

In accordance with the theoretical expectations of herd immunity, we found that both the *Sus* strain and the phage, were maintained in the population throughout the course of the experiment (Fig. 2D, E). This coexistence was essential for recombination to occur. When *Immu* was replaced by a strain carrying the same *yfp-kanR* cassette but lacking the phage targeting spacers, almost no recombination was observed (Suppl. text, Fig. S5). When the initial ratio of *Immu* to *Sus* was far from the herd immunity threshold, *Immu* failed to maintain *Sus* and phage in the population (modelled in Fig. S3, S4). This resulted in elimination of gene flow and HR (Fig. S5). Peak phage concentration can be observed at 3 hpi, when the phage initially had replicated on exponentially growing *Sus* cells, and after this point the phage concentration converged back to an equilibrium concentration around 10^6^ PFU/ml (Fig. 2E).

### Recombination is sustained in periodically resting populations

The number of recombinant CFUs kept increasing between 6 and 24 hpi (Fig. 2A). However, because phage P1 replication is inhibited in resting bacteria (*34*), recombination is expected to halt eventually. To determine the long-term population dynamics in stationary phase when recombination begins to fade out, we altered the experimental conditions such that stationary phase was reached earlier in the experiments. Again, we mixed overnight cultures of the two strains in a ratio 95% *Immu* to 5% *Sus*. Then, we inoculated phage P1 at initial MOI=0.1 and diluted the cultures 1:2 with fresh medium without selection antibiotics to allow bacteria to grow for ∼1 generation before the cultures reached stationary phase. We incubated the cultures for 24 hours and sampled at 3, 6, 12, 18 and 24 hpi onto selection agar plates to enumerate recombinant and mutant CFUs.

Under these conditions, the number of recombinant CFUs increased rapidly already at 3 hpi (Fig. 3A, solid line), significantly (t = 14.76, p-value < 0.001) exceeding the number of mutant CFUs (Fig. 3A, dashed line). This observation confirms that transduction-mediated recombination is indeed induced in early stationary phase. From 6 to 24 hpi we observed a slow decrease in the number of recombinant CFUs, with a slight increase between 18 and 24 hpi. This decrease was, however, partially caused by fluctuations in the total bacterial concentration (Fig. 3B), indicated by a relatively constant frequency of recombinants between 6 and 24 hpi (Fig. 3C). Similarly to the previous experiment, we observed a relatively stable fraction of phage-susceptible cells fluctuating around the herd immunity threshold concentration (Fig. 3D) and convergence of the number of phages to their equilibrium concentrations (Fig. 3E).

**Figure 3.**
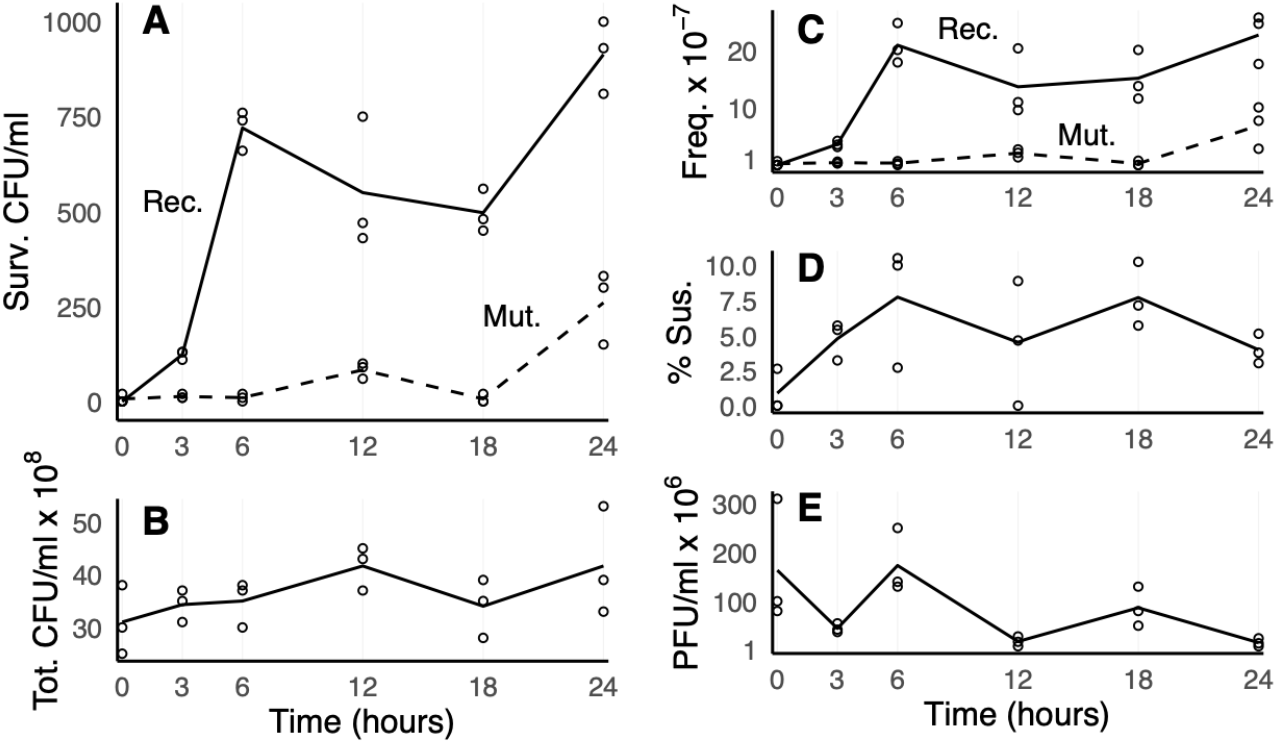
Recombination commences in early stationary phase. **(A)** In cultures, which were initially diluted only 1:2, the concentration of recombinants (solid line, Rec.) starts increasing at 3 hpi and halts after 6 hpi. **(B)** Total CFU concentration increases from initial 1:2 dilution and then fluctuates around stationary phase cell culture concentrations. **(C)** Frequency of recombinants increases between 0 and 6 hpi and then fluctuates around a constant value. **(D)** Phage-susceptible cells remain in cultures throughout the experiment. **(E)** Phage concentration slowly converges to its equilibrium concentration around 10^6^ PFU/ml. In all panels the shaded areas around the curves show standard error of the mean.

To find out whether recombination reinitiates after bacteria are given a burst of nutrients, we conducted another experiment, where we again inoculated the cultures 1:2 and at 12 and 24 hpi we diluted the cultures again 1:2. In Figure 4 we show that in the diluted cultures the number of recombinant CFUs surviving selection on plates significantly increased until the next sampling compared to both the previous sampling and the non-diluted controls. Since the samplings were always done before dilutions, thus in stationary phase, the concentration of recombinants reflected their frequency in the population (Fig. S2).

**Figure 4.**
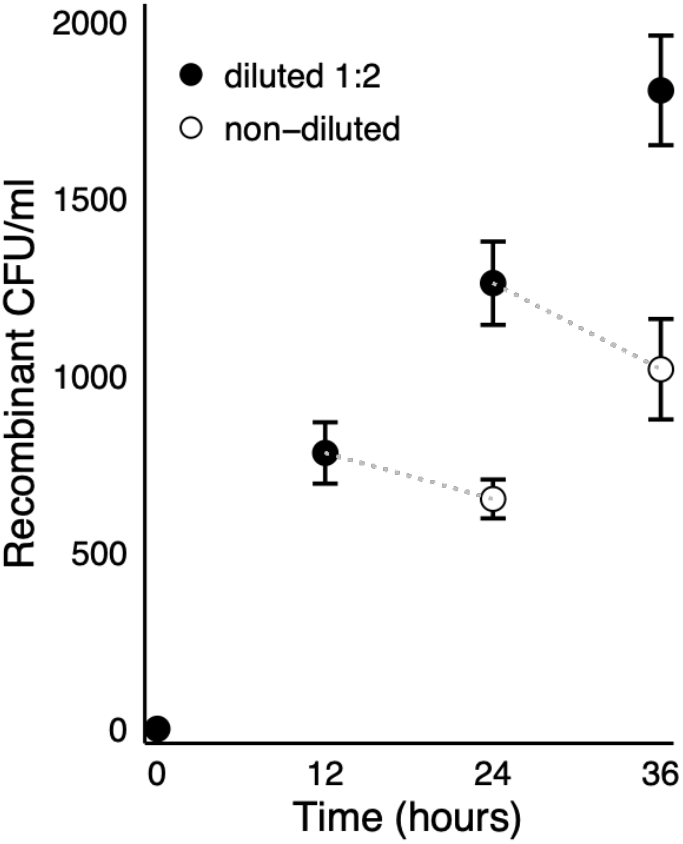
Recombination periodically recovers after short episodes of growth. The number of recombinant CFUs in cultures inoculated 1:2 increased significantly between 0 and 12 hpi (t = −9.01, p-value < 0.001), and then declined insignificantly until 24 hpi (empty circle, t = 1.26, p-value = 0.27). When cultures were diluted 1:2 at 12 hpi, the number of recombinants increased significantly at the sampling point 24 hpi (t = −3.28, p-value = 0.031), and again, without dilution declined insignificantly at the sampling point 36 hpi (empty circle, t = 1.21, p-value = 0.29). The number of recombinant CFUs in cultures diluted 1:2 12 hpi as well as 24 hpi again increased significantly when sampled at 36 hpi (t = −3.62, p-value = 0.022). Differences between mean recombinant CFU concentrations of diluted and non-diluted were significant both at 24 hpi (t = −4.69, p-value = 0.0094) and 36 hpi (t = −4.58, p-value = 0.01). Samples were always taken before dilution step i.e., from stationary phase saturated cultures. Error bars represent standard error of the mean.

The frequency of transduction in our experiments (Fig. 2F, 3C) was consistent with previous measurements, *i.e*., between 10^−5^ and 10^−6^ transductants per phage particle for P1 (*35*). Extrapolating from our experiments to the whole bacterial chromosome (all genes), up to roughly 1 in 1000 cells recombine a random sequence from the donor strain’s chromosome per generation (*i.e*., in each 1:2 dilution cycle), which from an evolutionary point of view, is a remarkably high recombination rate. In fact, such frequent HR can lead to random associations among mutations in many loci in just a few hundred generations!

### RecG mediates HR in stationary phase

The experiments described above show that cycles of nutrient depletion and nutrient influx favor HR within bacterial populations and imply that recombination is actively increased in stationary phase when nutrients become scarce.

To examine this hypothesis, we tested the known recombination regulatory pathways, which may play a role in recombination in stationary phase. We tested two main recombination initiation pathways: (i) *recFOR* by testing transduction efficiency in a *ΔrecF* mutant, and (ii) *recBCD* by testing *ΔrecB*. Furthermore, we tested two pathways that have been reported in the resolution of recombination intermediates: RuvABC, by testing a *ΔruvA* mutant and RecG, by including a *ΔrecG* mutant.

We measured transduction efficiency in the mutant strains transformed with our CRISPR immunity plasmid, grown to stationary phase. We also included strain *Immu* as a positive control. As a negative control we used *ΔrecA*, as *recA* is central to all HR pathways in bacteria (*36*).

As expected, recombination did not occur in Δ*recA* (Fig. 5). In Δ*recF*, we observed a two-fold reduction in the number of transductants compared to the wild type *Immu* strain, showing that the RecFOR pathway is only partially or indirectly involved in stationary-phase recombination. On the contrary, Δ*recB* exhibited no recombination at all, indicating that the molecular complex RecBCD mediates recombination events (*37*). The recombination intermediates formed by RecBCD/RecA that mediate strand exchange can be resolved through two distinct pathways, RuvABC and RecG (*38*). We tested both by using deletion mutants. In Δ*ruvA* we saw no significant difference from the wild-type *Immu* strain, indicating that the RuvABC resolvase complex is not involved. However, Δ*recG* showed 100% reduction in the number of transductants, thus complete hindrance of recombination, indicating that one of its main roles is to mediate HR in stationary phase.

**Figure 5.**
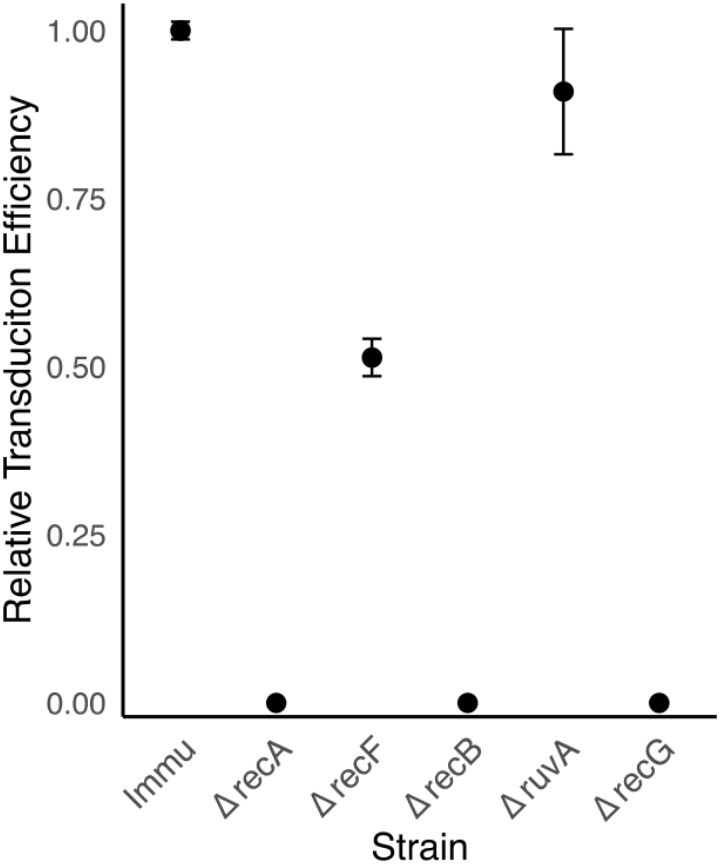
Transduction efficiency across deletion mutants. The wild-type *Immu* strain exhibits relative transduction efficiency = 1 (4147 recombinant CFU/ml per 5 x 10^8^ PFUs). *ΔrecA* shows no recombination as RecA is central to all recombination processes in the cell. *ΔrecF* (RecFOR pathway) shows a reduction in transduction efficiency to 51 % of the wild type *Immu* strain (t = 15.82, p-value < 0.001). In *ΔrecB* (RecBCD pathway) transduction efficiency drops to 0 (t = 75.6, p-value < 0.001). From two enzymes, which recruit RecBCD complex, *ΔruvA* and *ΔrecG*, the former shows no significant drop in transduction efficiency (t = 0.96, p-value = 0.39), whereas the latter shows transduction efficiency of 0. Error bars show standard error of the mean.

## Discussion

In this study, we showed that CRISPR immunity, in addition to providing defense against phage, enables frequent genome-wide HR. This process relies on protection of phage-susceptible bacteria by herd immunity, thus allowing a generalized transducing phage to establish gene flow from phage-susceptible to immune bacteria. Transduction-mediated recombination starts at the entry of stationary phase, when nutrients become scarce (*39*). When recombination peters out in late stationary phase, a moderate influx of nutrients, which briefly restores growth and subsequent re-entry into the stationary phase, suffices to reignite recombination. Since recombination occurs substantially more frequently than *de novo* beneficial mutations, not only it can drive adaptation by bringing beneficial mutations to the same genomic background, but it could also lead to high effective recombination rates (*r/m*) and help explain the near-random associations among mutations across loci in many bacterial species.

We identified *recG* to be essential for HR. RecG is a helicase, which is present in almost all bacterial species (*40*), and has been implicated to act at the replication fork end of a D-loop (*41*) and in the resolution of Holliday junctions in the absence of RuvABC (*42*). However, its involvement in many other cellular processes underscores that the role of RecG is still rather enigmatic (*43*). As *recG* is located within the *spo* operon, which is upregulated during bacterial stringent response, it has been suggested to be activated in response to starvation (*44, 45*). This view is consistent with regulation of RecG by the CreBC global regulatory system. CreBC is strongly upregulated during aerobic growth in minimal medium containing short-chain, low-energy carbon sources (formate, acetate, lactate, pyruvate, and malate), but its expression is suppressed on longer five- and six-carbon compounds (arabinose, fucose, gluconate, and glucose) and in nutrient broth, when cells generate energy by respiration on high energy glycolytic carbon sources. (*46, 47*). During growth in LB (used in this study), where amino acids are the main carbon source, rapid L-serine catabolism yields pyruvate, which peaks in late-exponential phase and is then used as a carbon source, leading to acetate production (*48, 49*). Therefore, accumulation of pyruvate and acetate, together with L-serine depletion, likely serves as an early indicator of impending nutrient scarcity, increasing *recG* expression via the CreBC regulon. A similar scenario can be expected during aerobic growth on other rich carbon sources, such as glucose and other mono- and disaccharides, which lead to overflow metabolism (the acetate switch) and acetate accumulation, and under anaerobic conditions (*48, 50*).

Taken together, the operon context and CreBC regulation suggest that *recG* boosts HR in response to impending resource scarcity, analogous to unicellular eukaryotes, which engage in sexual reproduction under starvation.

Herd immunity-driven HR has substantial implications for bacterial evolution. During exponential growth, the process of DNA replication generates *de novo* point mutations (*51*) and gene copy-number variants (*52*). As the environment transitions to resource scarcity signaling the onset of stationary phase, bacteria activate stringent response and slow down growth (*45*). Especially under starvation, increasing genotypic diversity can be advantageous, because it enables adaptation to newly available resources. However, resting bacteria are not known to generate *de novo* genetic variation for many days until long-term stationary phase, when GASP (growth advantage in stationary phase) mutants start replicating (*53*). Our population dynamics-based mechanism, which relies on host-phage coexistence and stationary-phase gene flow, bridges the gap between adaptation during exponential growth and adaptation in long-term stationary phase by reshuffling standing genetic variation.

CRISPR-Cas systems are primarily viewed as adaptive immune defenses against mobile genetic elements – mainly phages and conjugative plasmids – and thus as barriers to gene flow (*54*–*56*). They can also participate in other cellular processes, including regulation of gene expression, cell dormancy, DNA repair and genome evolution (reviewed in (*57, 58*)). One study reported that, in a laboratory assay, a CRISPR-Cas system can facilitate transduction in *Pectobacterium atrosepticum* (*29*). However, while those results suggest a potential role for CRISPR in facilitating HR, the experimental conditions suppressed any population-level dynamics and precluded inference about within-population recombination or evolutionary consequences: purified phage lysates were used at relatively high multiplicity of infection (MOI ∼ 1), with highly concentrated cells and multiple centrifugation/washing steps before plating. The authors explicitly argued that, in the presence of phages, CRISPR-resistant bacteria should outcompete susceptible bacteria, causing phage replication – and thus transduction – to cease. However, as we have shown here, this is not the case: through herd immunity, CRISPR actually helps preserve a reservoir of susceptible bacteria and thereby maintain genetic variability, which in turn enables gene flow capable of generating near-random associations among loci and high *r/m* ratios. Thus, our findings reveal a broader eco-evolutionary role that may help explain why these systems are so widespread (*32, 59*).

Several lines of evidence suggest that herd immunity-driven transduction-mediated HR is likely pervasive across natural populations. First, bacteria in the wild are outnumbered by phages about ten-fold (*60*), implying abundant potential transducing phage. For example, the vast majority (93.5%) of induced *S. enteric*a prophages were capable of generalized transduction (*61*). Comparative genomics further shows that recombined DNA block sizes match typical bacteriophage genome sizes, consistent with generalized transduction underlying widely detected HR (*15, 62*). In *Pseudomonas aeruginosa*, across 32 strains and 28 isolated phages, CRISPR–Cas systems targeted almost exclusively temperate phages (*63*), implying preferential targeting of generalized-transducing phages. Natural bacterial populations exhibit high diversity in CRISPR spacers (*64*), and in the presence/absence of various other immune systems (*65, 66*). This indicates that different fractions of the populations are immune to different phages – immune diversity potentially maintained by herd immunity – thereby enabling multidirectional, and potentially more frequent, gene flow.

As noted in the introduction, other bacterial immune systems, which interfere with phage replication at the DNA level (*e.g*., RM, BREX, DISARM (*67, 68*)), are also likely to give rise to herd immunity and may elevate gene flow through mechanisms similar to those presented here. Comparative genomic analyses have linked RM systems to elevated rates of gene gain and HR (*27, 28*). Generalized transduction can also be facilitated by lysogenic immunity, as lysogens can acquire transduced DNA from neighboring phage-susceptible cells in a process called autotransduction (*69*). Likewise, in many *Staphylococcus aureus* strains, plasmid transduction occurs to strains insensitive to the lytic action of the transducing phages due to protection by lysogenic immunity and RM systems (*70*).

Finally, the ecological conditions of natural bacterial populations are likely to support frequent, robust, herd immunity-driven HR. Doubling times in the wild are typically substantially longer than in the laboratory (*71*), and population sizes and nutrient availability often fluctuate (*72, 73*). This indicates that bacteria spend a substantial fraction of their life cycle transitioning between exponential and stationary phase – periods when recombination is more likely. Consistent with this, our mathematical model also indicates that such suboptimal growth conditions greatly expand the parameter space in which herd immunity can operate (Fig. S3, S4).

In conclusion, CRISPR is not only an efficient anti-phage defense system but can also turn phages – bacterial pathogens – into *de facto* pollinators that help bacteria adapt. Remarkably, this process is linked to the onset of resource scarcity and may help explain the widespread, yet puzzlingly high rates of HR observed across bacteria. Because most transducing phages have narrow host ranges, recombination is effectively confined to closely related bacterial strains, minimizing the risk of incorporating maladaptive or disruptive genetic material. Such confinement may act as a cohesive force, maintaining species integrity despite frequent genetic exchange (*74*). Conversely, ecological divergence – such as changes in phage receptor specificity – can alter phage host range and restrict gene flow, thereby reinforcing genetic isolation and potentially contributing to bacterial speciation (*16*–*18*).

This raises an overarching question: are CRISPR immunity and broader bacterial immune diversity maintained primarily for their individual defensive roles, or also because they enable a more general route to homologous recombination that substitutes for sexual reproduction in prokaryotes?

## Methods

### Bacterial strains construction

The *Immu* strain was a gift from Hande Acar Kirit and its construction is described in (*75*).In brief, it contains a chromosomal insertion of a cassette composed of a gene for yellow fluorescence protein (*yfp*, mVenus) under control of the constitutive phage lambda PR promoter, together with a gene for kanamycin resistance, at the phage P21 *attB* site of the parental MG1655 strain. This strain was transformed with the plasmid pCas9P1, which provided it with immunity to the P1 phage. pCas9P1 was derived from the pCas9 plasmid carrying a *Streptococcus pyogenes* truncated CRISPR type II system and chloramphenicol resistance. pCas9 was a gift from Luciano Marraffini (Addgene plasmid #42876). Two spacers targeting the P1 phage DNA were designed as oligonucleotides P1_DS_F and P1_DS_R (for all primer sequences see Table 1), annealed together and inserted into pCas9 following a protocol described in (*76*).

Next, a gene for red fluorescence protein (*rfp*, mCherry) was cloned into the AxyI restriction site downstream of a chloramphenicol resistance gene. mCherry with a T1 transcription terminator was amplified by PCR from pZS*21mCherry using primers AxyI_mCherry_F, and AxyI_T1_R, ligated into the pCas9P1 backbone and transformed into DH5alpha competent cells. It should be noted that our system lacks the genes for spacer acquisition because we aim to investigate the dynamics of an already resident immune and susceptible cells in ratios close to the herd immunity threshold frequency.

The *Sus* strain was constructed from the parental strain MG1655 by insertion of a cassette containing a gene for cyan fluorescence protein (*cfp*, mCerulean) under control of the constitutive promoter PN25, and a spectinomycin resistance gene. It was inserted into the phage λ *attB* site following a protocol described in (*76*). mCerulean was amplified by PCR from pTH2 (Addgene plasmid #84452) using a forward primer SphI_PN25_NdeI_CFP_F, which contained the PN25 promoter as an overhang, and a reverse primer HindIII_CFP_R. This strain was transformed with pCas9 containing no spacers, rendering the *Sus* strain chloramphenicol resistant but not immune to the P1 phage.

The P21 *attB* site containing the *yfp-kanR* cassette and the phage phage λ *attB* site containing the *cfp-specR* cassette are 389 kbp away from each other so that the two loci cannot be co-transduced by P1, which can only package up to 100 kb of DNA (*77*).

All constructs were sequence verified and confirmed by phenotypic testing for fluorescence and antibiotic resistance.

### Cultivation of bacteria and phages

Bacterial cultures were cultured in TPP round bottom 5 ml culture tubes (cat. #091106) in a New Brunswick Scientific Innova 43R Incubator Shaker at 37ºC and 250 rpm. LB broth (Sigma-Aldrich, cat. # L7275) was used as cultivation medium. LB broth with agar (Sigma-Aldrich cat. # L2897) was used to make LB agar plates.

All growth media and plates were supplemented with chloramphenicol 25 µg/ml. Liquid growth media were supplemented with 5 mM calcium chloride as it is required for adsorption of the phage P1 to bacterial cells. Plates used for selection of recombinant and *de novo* mutants (referred to as selection plates hereafter) were further supplemented with spectinomycin (35 µg/ml), kanamycin (35 µg/ml), and sodium citrate (10 mM) to suppress further phage adsorption on plates. Plates used for enumeration of colony forming units (CFUs) did not contain selective antibiotics (i.e. containing only chloramphenicol 25 g/ml and sodium citrate 10 mM). Plates for total CFU counts were incubated overnight at 37ºC, selection plates were incubated for 48h to make sure that even slowly growing mutant strains form visible colonies.

Phage used in the study was the virulent variant of the P1 phage (P1vir, referred to as P1 in the study) and was obtained from Calin Guet’s strain and phage collection. P1 phage lysate for recombination experiments was produced on laboratory wild-type E. coli K12 MG1655 strain by addition of stock phage lysate (produced on the same strain) into exponentially growing culture of E. coli K12 MG1655 (OD600 0.4-0.5) at MOI=1. Resulting lysate had a concentration 2.8 ± 0.2 * 10^10^ PFU/ml. Standard double-layer soft agar assays were used to enumerate the number of PFUs in cultures. Plates used for plaque assays were supplemented with chloramphenicol (25 µg/ml) and calcium chloride (5 mM).

### Competition assays

We inoculated *Immu* and *Sus* strains 1:1000 by serial dilution into fresh LB and combined the cultures in a 1:1 ratio in 2 ml culture tubes. At the time of inoculation, we sampled from the mixed culture in quadruplicates and plated serially diluted samples on LB plates supplemented with 25 μg/ml chloramphenicol. Cultures were incubated under shaking conditions as described above for 20 hours and sampled again.

### Recombination experiments

Exponential phase experiments were conducted as follows: overnight cultures were inoculated 1:100 in fresh LB medium supplemented as described above. P1 phage was added to cultures at MOI=0.1 and cultures were incubated as described above. At time intervals 3, 6, 12, 18, and 24 *hpi*, samples of 100 μl were spread onto selection plates. Simultaneously with sampling onto selection plates, samples were also serially diluted and plated onto LB agar plates without selective antibiotics (i.e. containing only 25 μg/ml chloramphenicol and 10 mM sodium citrate) to enumerate the total number of CFUs, and performed double-layer soft agar assays to enumerate the number of PFUs in the cultures. All experiments were done in triplicate.

### Plate imaging and colony enumeration

Plates were imaged using Zeiss AxioZoom.V16-Apotome2 fluorescent microscope equipped with 0.5x objective and filter cubes appropriate for CFP (Zeiss 47), YFP (AHF F46-003) and mRFP (Zeiss 63). The image of each plate was stitched from 12 individual images using Zeiss Zen Blue software. Colonies on plates were counted using OpenCFU 4.0.0+dfsg-2 colony counting software (*78*).

All colonies containing both, *yfp-kanR* and *cfp-specR*, were counted as recombinant CFUs no matter whether they also expressed *rfp*. About 1.5% of CFUs were not expressing *rfp*. We find two possible explanations for this observation. Because the phage cannot replicate on *Immu*, it cannot transduce the *yfp-kanR* cassette into *Sus*. Thus, the first possibility is that the nontargeting plasmid from *Sus* is transduced into *Immu* and by homologous recombination (HR) the *rfp* gene is replaced by a sequence, which is missing it, resulting in a plasmid conferring immunity but not expressing *rfp*. Then the *cfp-specR* marker can be transduced into this strain generating a recombinant colony expressing both *yfp* and *cfp* but not *rfp*. Second possibility, the plasmid not targeting the phage is transduced from *Sus* into an immune recombinant strain. However, considering the frequency of *Immu* in the population (Fig. 2B, 3C), we find this explanation less likely than the first one.

All colonies containing only one of the two cassettes, i.*e*., either *yfp-kanR* or *cfp-specR*, were counted as mutant CFUs.

### Knock-out strain experiments

Single gene knock-out strains from the Keio collection with knock-outs of *recA* (JW2669), *recB* (JW2788), *recF* (JW3677), *recG* (JW3627), and *ruvA* (JW1850) were transformed with pCas9P1 plasmid. Knock-out strains as well as the control *Immu* strain were grown overnight (for 18h) to their final OD, with the exception of *ΔrecA*, which grew substantially more slowly and reached its final OD after 42h of incubation, so it was inoculated 24h before all other strains. Samples from the cultures were serially diluted in SM buffer and plated on agar plates with 25 μg/ml chloramphenicol to enumerate the concentration of CFUs. 150 μl of each strain was mixed with 50 μl of phage lysate grown on *Sus* strain (*i.e*., containing spectinomycin resistance and *cfp* cassette) with a concentration 1 × 10^10^ PFU/ml, and incubated for 2 h in 37ºC in a shaking incubator. After incubation 100 μl of each culture was plated on selection plates and incubated for 48 h. Plates were subsequently imaged and colonies were calculated as described above. For details on CFUs, OD measurements and MOIs see Table S1. All experiments were done in triplicate.

### Statistical analyses and figures

Statistical significance of differences in data for Fig. 4 from different treatments and timepoints (diluted vs. non-diluted, 0 vs. 12, 12 vs. 24, 24 vs 36 hpi), for Fig. 5 (Immu vs. each deletion mutant), and for Fig. S1 (Immu vs. Sus at time 0 and 20 hpi) were tested by the following statistical methods. First, each pair of data was tested for equal variances using Levene’s test. Because the test did not reveal significant differences in variances in any of the sampled pairs, we used the two sample two-tailed Student’s *t*-test to compare whether the means of the samples differed significantly. For each pair of samples, *t*-statistics and *p*-value are reported. Since all experiments were performed in triplicates and the variances were equal, the number of degrees of freedom was equal to 4 for all statistics. All statistics were calculated using RStudio 2023.12.0+369.

Figures presenting data were generated using *RStudio 2023.12.0+369* and packages *ggplot2, readxl, dplyr, gridExtra, scales, patchwork, cowplot*, and *ggpubr*. Graphics were drawn using *Inkscape 1.3.2*.

## Acknowledgments

We thank Stuart Baird, Nick Barton, Max Joesch, Meriem El Karoui, Mato Lagator, Ivana Mašlaňová, Stepan Ovchinnikov, Roman Pantůček, Maroš Pleška, Jitka Polechová, and Bryan Wu for feedback on the manuscript. We acknowledge the core facility CELLIM supported by the Czech-BioImaging large RI project (LM2023050 funded by MEYS CR) for their support. This work was supported by: H2020 European Research Council EVOLHGT No. 648440 to JPB, European Commission Programme EXCELES project LX22NPO5103 “National Institute of Virology and Bacteriology” to PPl, and a bilateral Czech-Austrian mobility grant 8J3AT003 funded by MEYS CR and OeAD AT to PPa and CCG.

## Author contributions

Conceptualization: CCG, PPa

Methodology: CCG, PPa

Investigation: CCG, PPa

Mathematical modelling: PPa

Visualization: PPa

Funding acquisition: CCG, JPB, PPa, PPl

Project administration: PPa

Supervision: CCG, JPB, PPa

Writing – original draft: PPa

Writing – review & editing: CCG, PPa

## Supplementary Materials

Supplementary Information is available for this paper. PDF includes Supplementary Text, Fig. S1, Fig. S2, Fig. S3, Fig. S4, Fig. S5, Table S1, Table S2

Correspondence and requests for materials should be addressed to Calin C. Guet or Pavel Payne.

