## Supplementary text for "CRISPR herd immunity against a transducing phage underlies adaptive homologous recombination in resting bacteria"

### Supplementary Materials

#### Supplementary Text

##### *Efficiency of plating on Immu strain*

In the construction of the pCas9P1 plasmid we used two spacers to enhance immunity against the phage and to reduce the probability of phage escape mutants that can evade cleavage by CRISPR-Cas. The presence of multiple spacers targeting a single phage species is common in bacteria harboring CRISPR-Cas immunity and are acquired in a process called primed spacer acquisition (*1*). Efficiency of CRISPR immunity was determined using efficiency of plating (EoP) plaque assay on *Immu* relative to the *Sus* strain. EoP of *Immu* with two spacers was below the detectable limit, *i.e.*, less than $10^{-9}$.

##### *Competition assay of Immu and Sus strains*

To test whether *Immu* and *Sus* strains have comparable fitness, we performed exponential growth competition assays. In Figure S1A we show that at the time of inoculation (1:1000) in a 1:1 ratio, the concentrations of *Immu* and *Sus* did not differ significantly (t = -0.16222, df = 2.8594, p-value = 0.8819, mean diff.: 0.033 × $10^6$, 95% CI: -0.706 × $10^6$, 0.639 × $10^6$). After 20 hours of incubation (Fig. S1B) the mean concentration of *Immu* was slightly lower than that of *Sus*. However, the difference in means was not significant (t = -1.5492, df = 3.6697, p-value = 0.2025, mean diff.: 0.4 × $10^9$, 95% CI: -1.1430511 × $10^9$, 0.3430511 × $10^9$), indicating that ratios of the strains do not change substantially due to growth rate differences. It should be noted that the inoculation dilution in this experiment (1:1000) was 10x higher that in the recombination experiments (1:100). Thus, the slight and insignificant difference in growth rate is unlikely to affect the ratios of *Immu* and *Sus* in the experiments.

##### *Recombination rate inference*

From the stationary phase experiments (Fig. 3) with initial dilution 1:2 we infer that per 1 ml of culture, up to 689 recombination events of the *cfp-specR* reporter locus into the *Immu* background occur in a population with a concentration of 3.5 x $10^9$ CFU/ml. Because with 1:2 dilution each cell on average divided exactly once, this gives a frequency of 1.97 x $10^{-7}$ recombination events of the reporter locus per one generation. Considering the uniformity of the distribution of transduced blocks along the bacterial chromosome for phage P1 (*2*) and that *E. coli* K12 MG1655 contains 4,703 genes (*3*), we conclude that after one generation the frequency of recombinants of any two genes which cannot be co-transduced in the population is 9.26 x $10^{-4}$ CFU/ml. This gives an estimate that in this experiment, after one generation one in 1080 cells in the population undergoes recombination. It should be noted that we expect that recombination of the wild type allele lacking the *yfp-kanR* cassette replacing the *yfp-kanR* in *Immu* is occuring at the same rate as recombination the *cfp-specR* reporter locus into the *Immu* background. However, this does not double the overall rate of recombination because these events (just like recombination events of all other genes) remain undetected in our experimental setup.

***Herd immunity threshold estimation***

For calculation of the herd immunity threshold *H* we used a formula derived in (*4*)

$$H = \frac{\beta - 1 - \alpha\lambda}{\beta},$$

where $\beta$ represents the phage burst size, $\lambda$ its latent period and $\alpha$ the bacterial growth rate. We plugged in values for maximal P1*vir* burst size ($\beta$=85 pfu/cfu) and minimal latent period ($\lambda$=30 min) determined in (*5*), and a typical growth rate of *E. coli* in LB medium and 37ºC ($\alpha$=2.16) (*6*). This gave us the upper bound of the herd immunity threshold *H*=0.98 for nondepleted medium and exponentially growing cells. When we took into account reduction of burst size in depleted medium (*e.g.,* to 42% in medium depleted by 50% as in (*5*)), we arrive at *H*=0.95, which further drops when nutrients are further depleted until it becomes undefined when the phage reproduction is completely inhibited (*7*). Because it is difficult to estimate the average *H* in a culture, which initially grows and then reaches stationary phase, we operated with the ratio 5% *Sus* : 95% *Immu*. Such value is within the range of *H* for all experiments and allowed us to compare the results of individual experiments.

***Mathematical model of herd immunity***

In order to determine when herd immunity effectively protects susceptible cells and the phage in cultures growing initially exponentially and then transiting to stationary phase, we created a SIR (susceptible-infected-resistant) mathematical model of the system. Our model is described by the following set of four difference equations.

Immune (phage-resistant) cells $R_n$ grow by logistic growth according to an equation

$$R_n = \psi \cdot R_{n-1},$$

where $\psi$ describes the logistic growth term

$$\psi = 1 + r \cdot \left(1 - \frac{R_{n-1} + S_{n-1}}{K}\right).$$

Intrinsic growth rate $r$ is equal for both strains and carrying capacity $K$ is shared between immune and susceptible cells.

Susceptible cells grow according to the same formula

$$S_n = \psi \cdot S_{n-1} - S_{n-1} \cdot \frac{A \cdot P_{n-1}}{R_{n-1} + I_{n-1} + S_{n-1}},$$

but can also be removed from the system when they are infected by a phage. The phage has an adsorption coefficient $A$, which modifies the probability of phage adsorption on a susceptible cell and is defined as the fraction of phages that adsorb in current iteration. The infection probability is weighted by the concentration relative to phage-resistant, infected ($I_{n-1}$) and susceptible cells in the culture.

Infected cells are susceptible cells that have been infected by the phage and their concentration changes according to the following equation:

$$I_n = I_{n-1} + min\left(S_{n-1} \cdot \frac{A \cdot P_{n-1}}{R_{n-1} + I_{n-1} + S_{n-1}}, S_{n-1}\right) - H\left(I_{n-1}\left(1 - \frac{R_{n-1} + S_{n-1}}{\beta_0 K}\right)\right).$$

When phage concentration increases such that susceptible cells are infected by multiple phages, the number of newly infected cells is limited to the number of susceptible cells in the previous step, represented in the equation above by the min() extremum. Infected cells are removed from the system when they burst. The probability of bursting declines to zero when the bacteria reach $\beta_0$ fraction of the carrying capacity $K$, which is represented by the Heaviside step function $H$.

Finally, phage replicate on infected cells and are removed from the system when they adsorb to cells.

$$\begin{aligned} P_n = P_{n-1} &+ max\left(2 \cdot \beta \cdot \left(1 - \frac{I_{n-1} + S_{n-1}}{\beta_0 K}\right), 0\right) \\ &- R_{n-1} min\left(\frac{A \cdot P_{n-1}}{R_{n-1} + I_{n-1} + S_{n-1}}, P_{max}\right) \\ &- I_{n-1} min\left(\frac{A \cdot P_{n-1}}{R_{n-1} + I_{n-1} + S_{n-1}}, P_{max}\right) \\ &- S_{n-1} min\left(\frac{A \cdot P_{n-1}}{R_{n-1} + I_{n-1} + S_{n-1}}, P_{max}\right). \end{aligned}$$

Infected cell bursts with a per iteration burst size $\beta$, which is inversely proportional to the current growth rate of the cells and drops to 0 when bacteria reach $\beta_0$ fraction of the carrying capacity $K$. It should be noted that the parameter $\beta$ in the equation above is multiplied by the factor 2 because in our model, the phage replication is divided into two steps: First, susceptible cells are infected in one iteration, followed burst of these infected cells in the next iteration. Furthermore, we normalized the phage burst size as follows: The maximal reported P1*vir* burst size $\beta_{max} = 85$ requires the phage minimal latent $\lambda$=30 min (*5*), which is about 1.5 times longer than bacterial minimal doubling time (*6*). Thus, we divided the maximal burst size by the factor 1.5 to obtain the per iteration burst size $\beta = \frac{\beta_{max}}{1.5} = 57$.

Simultaneously, phages can be removed from the system when they adsorb to phage-resistant, infected, or susceptible cells with the adsorption coefficient $A$. The $P_{max}$ represents the upper limit of the number of phages that can adsorb to a single cell in one iteration.

Unless indicated otherwise, for all simulations we used the following parameters:

$P_0 = 1/10$ of total bacterial inoculum (MOI = 0.1),
$r = 1$,
$K = 10^9$,
$A = 0.9$,
$\beta = 57$,
$\beta_0 = 0.9$,
$P_{max} = 100$.

$R_n$ Number of phage-resistant cells in $n$-th iteration
$I_n$ Number of infected susceptible cells in $n$-th iteration
$S_n$ Susceptible cells
$P_n$ Phages
$r$ Intrinsic growth rate of immune and susceptible cells
$K$ Bacterial carrying capacity
$A$ Phage adsorption coefficient
$\beta$ Phage burst size
$\beta_0$ Phage fade out: A fraction of carrying capacity $K$ when phage stops replicating
$P_{max}$ Maximum number of phages which can adsorb to an immune cell (determines the rate of phage decay)

***Conditions for herd immunity***

Using our mathematical model, we conducted numerical simulations to determine the parameter range, where herd immunity is effective and how it depends on variable initial dilution factors of the cultures and phage burst size $\beta$. With high dilution factors the bacterial populations grow longer in exponential phase, which supports high burst size of the phage. If the initial ratio of *Immu:Sus* in the culture is low, the phage replicates on the susceptible cells too fast and the sudden phage burst kills all *Sus* cells before *Immu* can protect them through herd immunity (Fig. S3, S4A). Similarly, the effectivity of herd immunity depends on phage burst size $\beta$ (Fig. S3, S4B). With high $\beta$, the initial ratio must be close to the herd immunity threshold, whereas when $\beta$ is lower (typically in a suboptimal growth medium), much larger range of initial ratios is tolerated. When cultures reach polymorphic equilibrium in stationary phase, the free phage is depleted but it remains protected within nonreplicating infected cells.

***Test for essentiality of herd immunity***

Our modelling results indicate that herd immunity is essential for recombination to operate. To test this assumption, we conducted an experiment where we followed the same methodology as for the recombination experiments (Fig. 2 and 3) except we started with varying initial ratios of *Immu:Sus* and sampled only at 24 hpi.

In Figure S5 we show that when the initial ratio of *Immu:Sus* fell outside the parameter space where herd immunity supports coexistence (Fig. S3, S4), *i.e.,* 50% *Immu* to 50% *Sus*., the system did not lead to gene flow and recombination, most likely because of a rapid lysis of all susceptible cells. With initial ratios 90%:10% and 95%:5% we observed high frequency of recombination, as predicted by the model. The number of recombinants was highest with initial ratio 90%:10% likely because it is further from the equilibrium frequency of herd immunity threshold, thus during the exponential growth a majority of susceptible cells lyse as simulated in Fig. S3, and provide more copies of the *cfp-specR* cassette for transduction.

With the initial ratio 99%:1%, we observed on average only 10 recombinant CFU/ml, indicating that when the system deviates from the herd immunity threshold frequency of *Immu* towards fixation, the *Sus* concentration is insufficient to provide enough donor genetic material for recombination. However, in a situation where immunity induces a cost, the frequency of *Sus* should eventually converge to 1-*H*.

Furthermore, we tested the conditions under which both the *yfp-kanR* strain and the *cfp-specR* strained carried the susceptible plasmid pCas9 without P1*vir* targeting spacers. In these conditions herd immunity could not operate, the cultures lysed and did not produce nearly any recombinants.

These results confirm that herd immunity is essential for coexistence of susceptible and immune strains and for stable and frequent recombination within the populations.

***Knock-out strain experiments***

Detailed results including incubation time, optical density at OD600 readings (in 1:10 virtual dilution), CFU concentrations, PFU/CFU ratios, and concentrations of recombinant CFUs for each strain are shown in Table S2.

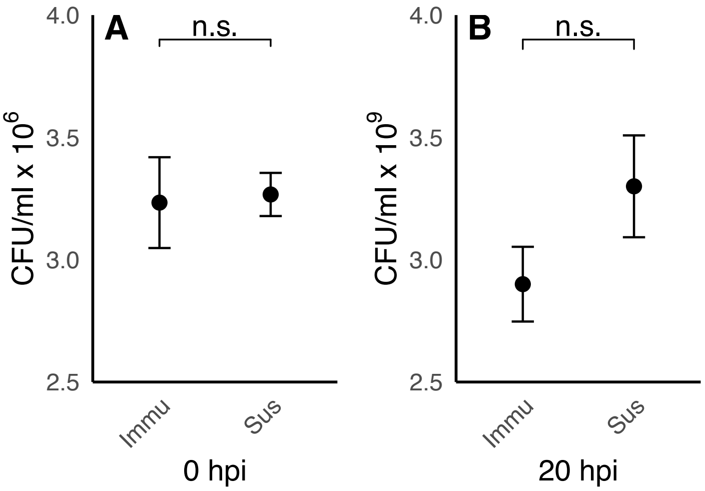


**Figure S1. Concentration of CFU in competition experiment.** (**A**) Concentrations of CFUs of *Immu* and *Sus* did not differ significantly right after inoculation 1:1000 ($t = -0.16$, p-value = 0.88). (**B**) Concentration of *Immu* was slightly lower than of *Sus* at sampling 20 hpi but the difference was not significant ($t = -2.08$, p-value = 0.11 ).

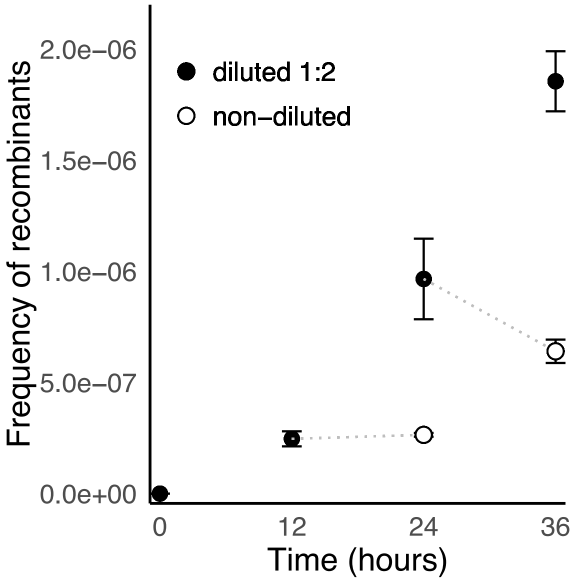


**Figure S2. Frequency of recombinants in sequentially diluted cultures.** Frequency of recombinant CFUs in cultures inoculated 1:2 increased between 0 and 12 hpi and then remained constnat until 24 hpi (empty circle at 24 hpi). When cultures were diluted 1:2 at 24 hpi, frequency of recombinants increased again (full circle) at the sampling point 36 hpi, and without dilution declined slightly (empty circle). Samples were always taken before dilution, thus sampling from a population in stationary phase with a comparable concentration of bacteria.

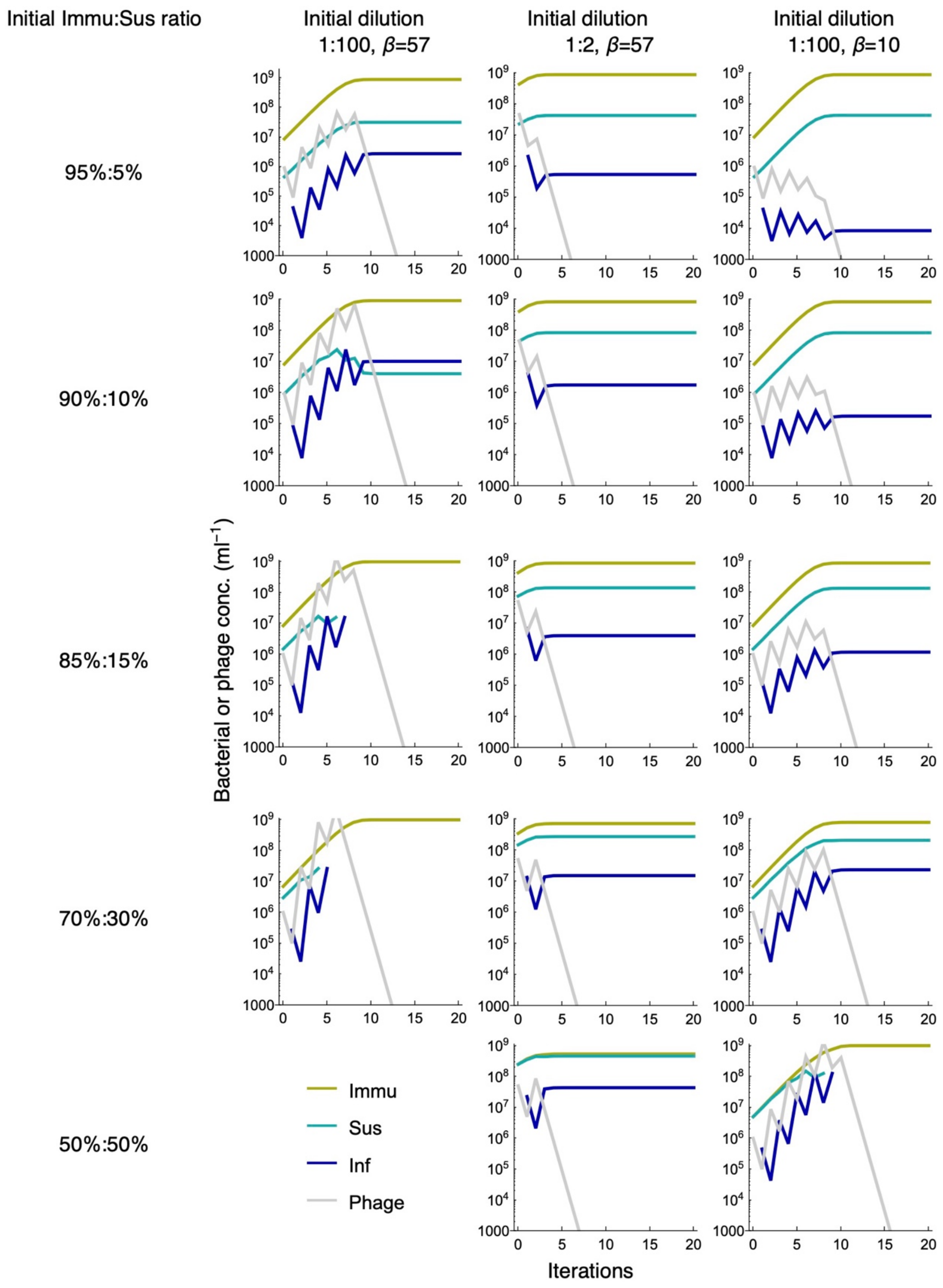


**Figure S3. Numerical simulations with selected parameters**. With initial dilution 1:100 (first column) the initial ratio of *Immu*(yellow):*Sus*(cyan) cells allowing herd immunity needs to be close to the herd immunity threshold. With initial dilution 1:2 (second column), herd immunity is effective for all selected ratios. With initial dilution 1:100 and burst size $\boldsymbol{\beta} = \mathbf{10}$, herd immunity is maintained down to 70%:30% but fails at 50%:50%. When cultures reach equilibrium in stationary phase, the free phage is depleted (gray curves) but it remains protected within nonreplicating infected cells (blue curves).

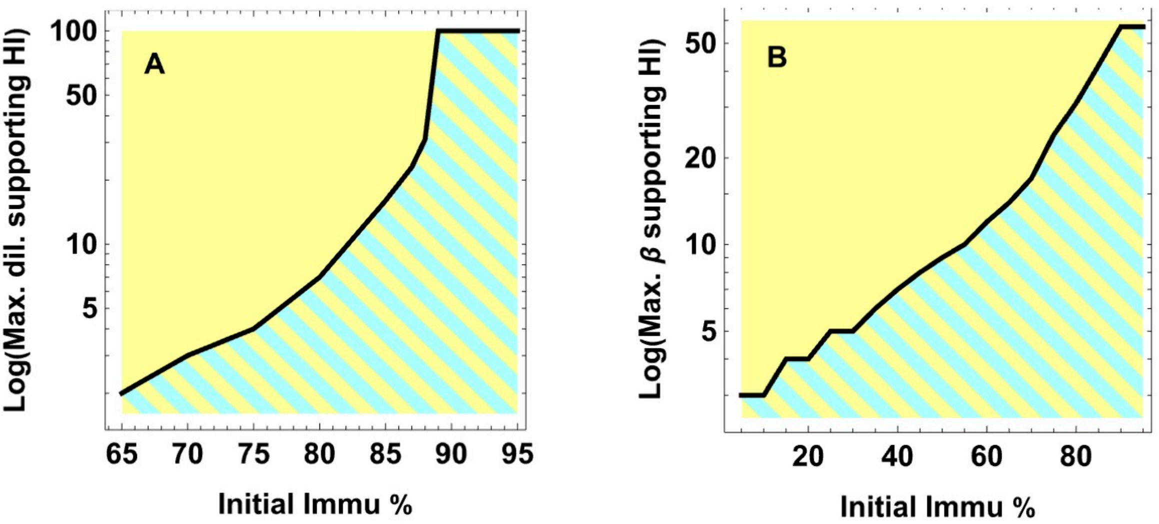


**Figure S4. Numerical analysis of conditions when herd immunity protects susceptible strain**. In the areas above the curves (yellow areas), only the phage-immune strain persists in the populations. Under the curves (cyan and yellow striped areas), both strains are maintained in the populations by herd immunity. (**A**) Maximal dilution factor (y-axis) still supports herd immunity for varying initial ratios of *Immu:Sus* (x-axis). Low dilution factors allow herd immunity to protect susceptible cells under larger range of initial *Immu* %. (**B**) Maximal burst size $\boldsymbol{\beta}$ (y-axis) still supports herd immunity for varying of *Immu:Sus* (x-axis). Lower burst size allows herd immunity to operate in larger range of initial *Immu* %.

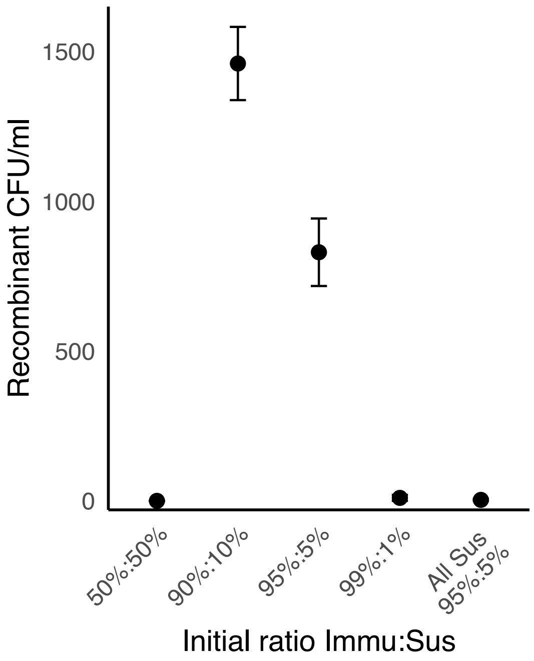


**Figure S5. Recombinants with varying initial ratios of strains.** With initial ratio at 50% *Immu*:50% *Sus* we observed no recombinants, whereas with 90%:10% and 95%:5% we see high frequency of recombination, which decreases to only 10 CFU/ml with 99%:1% ratio. When none of the strains carried the CRISPR immunity providing plasmid, cultures lysed and produced almost no recombinants (0.33 CFU/ml).

| Primer | Sequence 5' => 3' |
|---|---|
| P1_DS_F | AAAACAATAAGTGTGTGCTGGAGGGAAACCGCATTAGTTTTGGGACCATTCAAAACAGCATAGCTCTAAAACAAAAAGCAGAGAAGTATCCGTACTTTGAGG |
| P1_DS_R | AACCCTCAAAGTACGGATACTTCTCTGCTTTTTGTTTTAGAGCTATGCTGTTTTGAATGGTCCCAAAACTAATGCGGTTTCCCTCCAGCACACACTTATTG |
| AxyI_mCherry_F | ATGCCTGAGGGAGAGGAGTTATGGTTTCCAAGGGCGAGGA |
| AxyI_T1_R | GCGCTCGGGGATAAAACGAAAGGCCCAGTCT |
| SphI_PN25_NdeI_CFP_F | GCGCGCATGCTCATAAAAAATTTATTTGCTTTCAGGAAAATTTTTCTGTATAATAGATTCATAAATTTGAGAGAAGAGTTTACATATGAGTAAAGGAGAAGAACTTTTCAC |
| HindIII_CFP_R | GCGCAAGCTTTTACTTGTACAGCTCGTCCAT |

**Table S1. Primers used for molecular cloning in the study.**

| Strain | Incubation time (h) | OD600 (1:10) | CFU/ml | MOI (PFU/CFU) | Recombinant CFU/ml |
|---|---|---|---|---|---|
| *Immu* | 18 | 0.55 | $5.4 \times 10^9$ | 0.62 | 4240, 4050, 4150 |
| Δ*recA* | 42 | 0.16 | $9.7 \times 10^8$ | 3.44 | 0, 0, 0 |
| Δ*recB* | 18 | 0.39 | $3.0 \times 10^9$ | 0.9 | 0, 0, 0 |
| Δ*recF* | 18 | 0.51 | $4.0 \times 10^9$ | 0.83 | 1900, 2250, 2240 |
| Δ*recG* | 18 | 0.45 | $3.3 \times 10^9$ | 1.01 | 0, 0, 0 |
| Δ*ruvA* | 18 | 0.43 | $3.4 \times 10^9$ | 0.98 | 4220, 3000, 4090 |

**Table S2. Transduction experiments with knockout strains.** Details on each strain used in the experiments with knockout strains.
